# Cortical over-representation of phonetic onsets of ignored speech in hearing impaired individuals

**DOI:** 10.1101/2023.06.26.546549

**Authors:** Sara Carta, Emina Aličković, Johannes Zaar, Alejandro López Valdes, Giovanni M. Di Liberto

## Abstract

Hearing impairment alters the sound input received by the human auditory system, reducing speech comprehension in noisy multi-talker auditory scenes. Despite such challenges, attentional modulation on the envelope tracking in multi-talker scenarios is comparable between normal hearing (NH) and hearing impaired (HI) participants, with previous research suggesting an over-representation of the speech envelopes in HI individuals (see, e.g., Fuglsang et al. 2020 and Presacco et al. 2019), even though HI participants reported difficulties in performing the task. This result raises an important question: What speech-processing stage could reflect the difficulty in attentional selection, if not envelope tracking? Here, we use scalp electroencephalography (EEG) to test the hypothesis that such difficulties are underpinned by an over-representation of phonological-level information of the ignored speech sounds. To do so, we carried out a re-analysis of an EEG dataset where EEG signals were recorded as HI participants fitted with hearing aids attended to one speaker (target) while ignoring a competing speaker (masker) and spatialised multi-talker background noise. Multivariate temporal response function analyses revealed that EEG signals reflect stronger phonetic-feature encoding for target than masker speech streams. Interestingly, robust EEG encoding of phoneme onsets emerged for both target and masker streams, in contrast with previous work on NH participants and in line with our hypothesis of an over-representation of the masker. Stronger phoneme-onset encoding emerged for the masker, pointing to a possible neural basis for the higher distractibility experienced by HI individuals.

**Significance Statement:** This study investigated the neural underpinnings of attentional selection in multi-talker scenarios in hearing-impaired participants. The impact of attentional selection on phonological encoding was assessed with electroencephalography (EEG) in an immersive multi-talker scenario. EEG signals encoded the phonetic features of the target (attended) speech more strongly than those of the masker (ignored) speech; but interestingly, they encoded the phoneme onsets of both target and masker speech. This suggests that the cortex of hearing-impaired individuals may over-represent higher-level features of ignored speech sounds, which could contribute to their higher distractibility in noisy environments. These findings provide insight into the neural mechanisms underlying speech comprehension in hearing-impaired individuals and could inform the development of novel approaches to improve speech perception in noisy environments.

## Introduction

In multi-talker scenarios, comprehension of a selected speech stream (target) is hampered by the presence of competing speech streams (maskers). Our auditory system avails of spectro-temporal and spatial cues to segregate the target from the masker streams. Nonetheless, masker sounds cannot be fully ignored, as that would prevent us from switching our focus of attention at will. In fact, a wealth of research on the segregation of target and masker sound streams in the human cortex shows that neural signals exhibit an encoding of a combination of sounds, not only the target stream (Ding and Simon, 2012; Mesgarani and Chang, 2012; O’Sullivan et al., 2015b). Research on the neurophysiology of attentional selection sheds light on this process in normal hearing (NH) participants with both invasive and non-invasive neural recordings, indicating that the focus of attention modulates the neural processing of progressively more abstract speech properties, from the sound envelope, to phonemes, and lexical information (O’Sullivan et al., 2015b; Broderick et al., 2018; O’Sullivan et al., 2019; Di Liberto et al., 2021; Teoh et al., 2022). However, there remains considerable uncertainty on how attentional selection unfolds in hearing impaired (HI) individuals. This is particularly important to understand their comprehension difficulties in noisy multi-talker scenarios, leading to high distractibility and an increased listening effort (Ohlenforst et al., 2017). Here, we test the hypothesis that the cortex of HI individuals exhibits an over-representation of the masker speech stream in challenging listening scenarios (Lunner et al., 2020).

Acoustic properties of masker speech streams, such as the sound envelope, have been shown to be reliably encoded in neural activity (Brodbeck et al., 2020). However, it was only recently that neurophysiology studies could analyse the cortical encoding of acoustic and phonological speech properties in realistic multi-talker scenarios (Teoh et al., 2022). Specifically, EEG signals recorded in young NH individuals exhibited robust envelope and phonetic-feature encoding for the target speech, with weaker envelope tracking and no significant phonetic feature encoding for the masker speech. While previous work suggested a similar impact of attentional modulation on envelope tracking for both NH and HI participants, even with a certain degree of envelope over-representation with hearing difficulties (Fuglsang et al., 2020), the impact of attentional modulation on the neural encoding of phonetic features in HI individuals remains to be defined.

Investigating this issue is critical to understand the hearing and attentional strategies employed by HI listeners in naturalistic multi-talker environments (Alickovic et al., 2020; Lunner et al., 2020; Seifi Ala et al., 2020; Andersen et al., 2021; Fiedler et al., 2021; Shahsavari Baboukani et al., 2022). Previous studies have presented us with a complex array of findings as to how the degree of hearing loss changes how attentional modulation influences the disparity in the neural tracking of target and masker speech streams (Petersen et al., 2017; Fuglsang et al., 2020). As with NH listeners, higher-order speech processing was suggested to more greatly reflect attentional focus (Alickovic et al., 2021; Andersen et al., 2021), with increased top-down active suppression of the masker speech under more challenging listening conditions (Fiedler et al., 2019). However, such results relied on the temporal latencies of the neural responses – with longer latencies representing higher-order processes (Power et al., 2012; Puvvada and Simon, 2017) – without pinpointing the exact acoustic-linguistic properties that are encoded and suppressed in HI individuals.

Here, we investigated the neural encoding of target and masker speech streams in HI individuals, with the hypotheses that: 1) HI individuals would exhibit stronger cortical encoding of phonetic features for the target than for the masker speech; and 2) the encoding of some phonetic feature properties would be over-represented in HI individuals, potentially explaining the listening difficulties and high distractibility of HI individuals in multi-talker auditory scenes (Orf et al., 2023). To test these hypotheses, we carried out a re-analysis of an existing EEG dataset (Alickovic et al., 2021), where scalp EEG signals were recorded from HI individuals as they performed a selective-attention task in an realistic multi-talker scene. Multivariate regression was used to estimate the Temporal Response Function (TRF) describing the linear mapping from acoustic and phonetic features to the EEG signals (Crosse et al., 2016; Crosse et al., 2021). TRF models corresponding to target and masker speech streams were studied, providing us with objective neural measures of the cortical encoding of acoustic and phonetic features for testing the hypotheses of this study. Our results complement the existing literature, offering novel insights on the impact of attentional selection on the neurophysiology of speech in HI individuals, and contributing to future hearing impairment research and hearing-aid development.

## Methods

### Subjects

Thirty-four subjects (10 females, aged between 21 and 84 years, mean age = 64.2 years, standard deviation = 13.6 years) gave their written informed consent to participate in this study. All participants were native Danish speakers and had mild-to-moderately severe symmetrical sensorineural hearing impairment. They were experienced hearing-aid users, and were fitted binaurally with two hearing aids for this experiment. Participants reported no history of neurological or psychiatric disorders and had normal or corrected-to-normal vision. For further details on their hearing profile, hearing-aid fitting, and signal processing, see (Alickovic et al., 2021).

### Experimental design

Participants were presented with speech stimuli in a free-field cocktail-party scenario. Speech stimuli were delivered as participants were comfortably seated inside a sound-proof booth, at the centre of a circle of six loudspeakers, placed at ±30°, ±112.5° and ±157.5° azimuth relative to their location (**Figure 1A**). The task consisted of listening to a particular speech stream, corresponding to a specific talker and spatial location, while ignoring all other talkers. The target talker always originated from one of the two frontal loudspeakers (*S1* and *S2*), while the other frontal loudspeaker had to be ignored (masker). Both target and masker were presented at 73 dB sound pressure level (SPL). The other four loudspeakers in the background (*B1*-*B4*) were simultaneously presenting a 4-talker babble noise each, at a level of 64 dB SPL each. This created a 16-talker noise babble in the background. Audio stimuli were presented at a sampling rate of 44100 Hz, delivered through a sound card (RME Hammerfall DSP multiface II, Audio AG, Haimhausen, Germany) and played through six loudspeakers Genelec 8040A (Genelec Oy, Iisalmi, Finland). EEG data was simultaneously acquired with a sampling rate of 1024 Hz, from sixty-four electrodes mounted according to the International 10/20 system and two reference electrodes placed on the mastoids, using a BioSemi ActiveTwo system.

**Figure 1.**
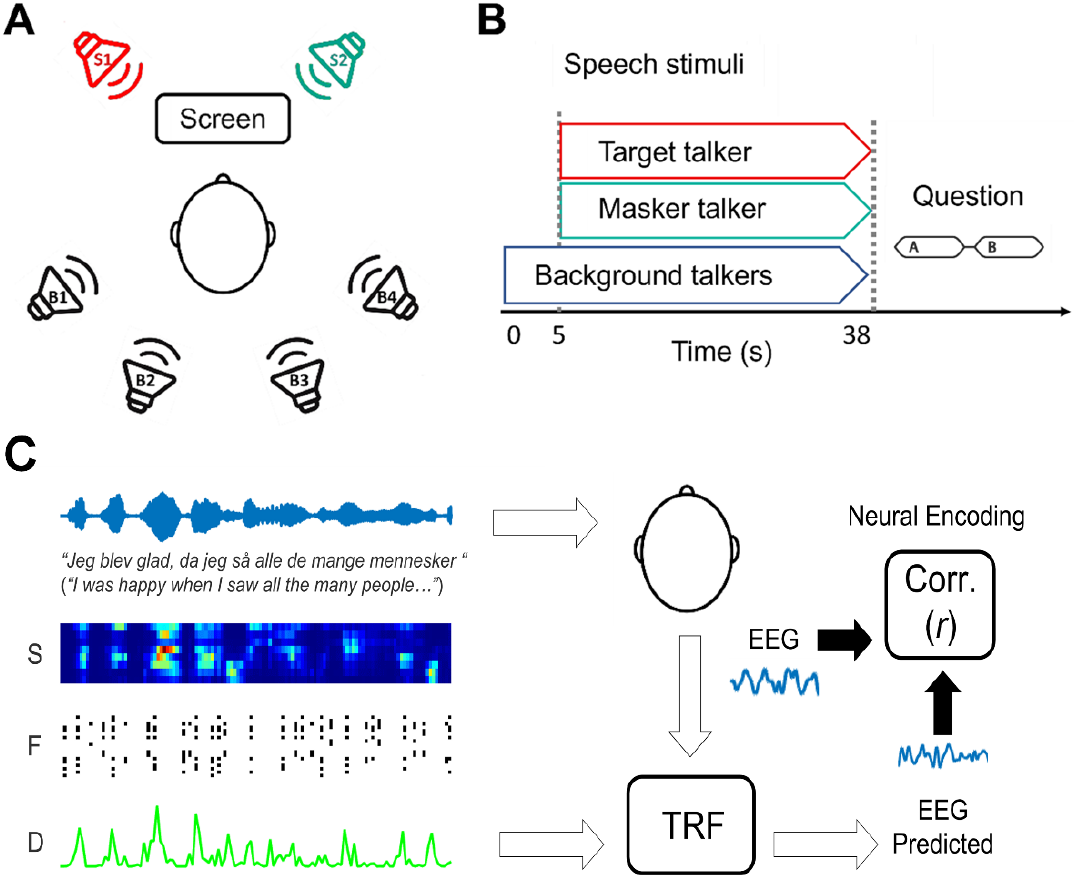
Experimental design and analysis framework. **(A)** Location of loudspeakers (for this particular example, S1: target speech, S2: masker speech, B1-B4: babble noise) relative to the participant. **(B)** Timeline of each experimental trial. Start of the noise babble at t_0_ = 0, followed by target and masker at t_1_ = 5s. **(C)** Neural signals were recorded with EEG as participants listened to natural speech monologues. A lagged linear regression analysis was carried out to estimate the Temporal Response Function (TRF) describing the relationship between speech phonetic features and low-frequency EEG (1-8 Hz).

This study presents a novel re-analysis of a previously existing dataset from Aličković et al. (2020, 2021). Note that only two out of the four experimental blocks were re-analysed as the rest of the data was not relevant to the goals of this study. As such, the results were derived based on 40 trials (20 trials per block).

The 2 blocks analysed here involved the identical selective-attention task, with the difference that the participants’ hearing aids were either utilizing a NR scheme or not, i.e., the NR ON and NR OFF conditions, respectively. In the NR ON block, the NR scheme used a minimum-variance distortionless response beamformer to attenuate the babble noise coming from the background (Kjems and Jensen, 2012). In the NR OFF block, the NR scheme was deactivated and, as such, participants were completely exposed to the background noise. Each block (20 trials) was composed of 4 sub-blocks of 5 randomized consecutive trials for each of “left male (LM),” “right male (RM),” “left female (LF),” and “right female (RF).” Before each sub-block of 5 trials, a visual cue was presented on the screen located in front of the participants, indicating the gender and location of the frontal speaker they needed to attend to. Participants underwent a familiarisation phase, with a brief training of 4 trials, one for each of the above combinations of target speakers’ sex and location of presentation. Each trial started with the presentation of the background babble noise and, after 5 seconds, the frontal target and masker speakers were added to the auditory scene (**Figure 1**). Following the end of the trial, participants were required to answer a two-alternative-forced-choice question on the target-speech content to check for their sustained attention to the task. After each block, participants could take a self-paced break before continuing with the experiment.

Individual speech stimuli consisted of short Danish newsclip monologues of neutral content. To control for differences between male and female target talkers, stimuli were all root-mean-squared (RMS) normalized to the same overall intensity. All silences were shortened to 200 ms and the long-term average spectrum of the babble noise was matched to the overall spectrum of male and female foreground talkers to avoid inconsistencies in masking.

### Stimuli and feature extraction

Here, we sought to determine how specific speech properties in this competing-talker attention scenario contribute to the neural-tracking response of target and masker speakers. Therefore, the foreground speech streams were modelled according to a set of features representing their low-level acoustic and phonetic characteristics. For the former set of features, we first extracted the stimuli’s spectrograms by applying a filter which partitioned the sound waveforms into eight logarithmically spaced frequency bands, from 250 Hz to 8 kHz (Hamming window, with a length of 18ms, a shift of 1ms; FFT with 1024 samples), according to Greenwood’s equation (Greenwood, 1961). We also extracted, for each frequency band, its half-wave rectified spectrogram derivative. Together, these acoustic features are referred to as ‘**S**.’

A second set of features was derived to capture categorical phonetic features, describing the sounds typical of the Danish language in terms of articulatory features, e.g., voicing, manner of articulation, and place of articulation. This feature-set is referred to as ‘**F**.’ First, speech stimuli were automatically transcribed, and the accuracy of the transcription was verified by a native Danish speaker. The transcripts were then processed by NordFA, a forced phonetic aligner for Nordic languages (Rosenfelder et al., 2014; Young and McGarrah, 2021), which identified timestamps corresponding to the start and end of each word into its constituent phonemes. The quality of the forced alignment was manually verified using Praat software on about 10% of the speech material, and with custom-made MATLAB scripts to assess that phoneme onsets corresponded to increases in the envelope, as expected in case of satisfactory phoneme alignment. Each phoneme was then represented as a binary vector with eighteen dimensions, each corresponding to one articulatory feature. Specifically, the following eighteen features were considered: syllabic, long, consonantal, sonorant, continuant, delayed-release, approximant, nasal, labial, round, coronal, anterior, dorsal, high, low, front, back, tense.

### EEG data preprocessing

Neural data were analysed by using MATLAB software (MathWorks). Analysis code was customized based on publicly available resources (please see the CNSP initiative website: https://cnspworkshop.net). EEG data were bandpass filtered between 1 and 8 Hz (Zion Golumbic et al., 2013; Di Liberto et al., 2021) using a zero-phase shift 4^th^ order Butterworth filter and then downsampled from 1024 to 64 Hz. Noisy EEG channels with a variance three times greater than that of the surrounding sites were replaced by means of a spherical spline interpolation, using a library in the EEGLAB Software. Subsequently, we re-referenced EEG signals to the average of the two mastoid channels. Single-subject EEG was also standardised by its overall standard deviation, preserving the ratio across electrodes.

### The Temporal Response Function (TRF) framework

We employed a system identification technique, known as the TRF framework, to estimate the relationship between neural signals and speech features, which can be conventionally approximated as a linear mapping. The TRF represents the optimal set of weights, obtained by employing regularized linear regression (Crosse et al., 2016), describing the transformation from the stimulus features to the corresponding brain responses. To avoid overfitting, leave-one-out cross-validation was applied across trials, by exploring a parameter space spanning from 10^−6^ to 10^4^, in search for the optimal lambda, which was selected as the regularisation value yielding the highest Pearson’s correlation between predicted and observed EEG data (forward model). Please note that in the case of multivariate speech representations consisting of both categorical and continuous variables (e.g., the FS model with spectrogram, spectrogram derivative and phonetic features) a banded regression was applied, in which the lambda search allowed for a different lambda selection for each set of features, aiming at the optimal combination of lambda values. Considering that the relationship between any stimulus feature and the resulting EEG response is not instantaneous, TRFs are always estimated by taking into account multiple time-lags. In this case, we considered a time-lag window spanning the interval from -100 ms to 350 ms. Since the stimulus-EEG relationship is estimated for each electrode separately and defined by examining the stimulus effect on the EEG at multiple latencies, TRF weights are interpretable both spatially, through topographies representing the scalp locations where this relationship is stronger or weaker, and temporally, by examining how the stimulus’ impact on the EEG signal evolves over time.

### Speech feature models

The TRF framework was used to determine the relationship between low-frequency EEG signals and the phonetic information in speech. For both NR-ON and NR-OFF conditions, single-subject TRF models were fit for target and masker stimuli separately.

First, forward TRF models were fit considering the phonetic features (F feature-set), yielding the F model, and a control F_*sh*_ model, with shuffled phonetic features (shuffled F feature-set). Comparing the predictive performance of the F and the F_*sh*_ models allows us to isolate neural variance that can be explained by the phoneme identities but not their timing alone (as F*sh* only contain phoneme onset information). Next, TRFs were also fit by considering low-level acoustic features (S feature-set), and a combination of F and S (FS feature-set combination). Here, we subtracted the EEG prediction correlations for the FS and S models deriving a metric called PhCat (also called FS-S in the literature), which quantifies EEG variance explained by phonetic features but not by the acoustic features i.e., acoustically invariant responses. In case of a significant predictive gain due to the addition of phonemes in the PhCat metric, we planned to run additional control analyses to determine whether the effect was due to the encoding of the identity of phonetic feature categories or, instead, to the encoding of their timing i.e., phoneme onsets (please note that the acoustic onsets are already accounted for by the spectrogram derivative contained in S). To do so, the TRF model fit was re-evaluated after a random shuffling of the phonetic features identity, producing a metric defined as PhOnset = F_*sh*_S-S

### Phonetic distance maps

To determine how attentional selection alters the sensitivity to phonetic features, EEG phoneme-distance maps (PDM; Di Liberto et al., 2015; Di Liberto et al., 2021) were compared for the target and masker TRF weights of the PhCat metric (FS-S). The rationale is to project the TRF weights onto a multi-dimensional space where the Euclidean distance between phoneme pairs corresponds to the difference in their EEG responses. In doing so, these maps allow us to determine the sensitivity of the EEG signals to specific phonetic features. PDMs were obtained as follows: First, phoneme weights were calculated as a linear combination of the weights for the corresponding phonetic features. Then, a multi-dimensional scaling analysis (MDS) was carried out to reduce the dimensionality of the TRF weights, considering all time-lags and electrodes simultaneously, while preserving the standardized Euclidean distances between the EEG signals corresponding to different phonemes. The masker PDM was then mapped onto the reference target PDM, for each NR, by isomorphic scaling via Procrustes analysis (ref; MATLAB function procrustes).

Four sets of phonetic categories were selected arbitrarily to reflect the major phonetic articulatory groups, ensuring that phonemes belonging to each set would create perceptually relevant phonetic contrasts defining the phonetic features: manner of articulation, place of articulation (lips), place of articulation (tongue), and voicing. For example, when considering the place of articulation (tongue) phonetic set, it is possible to explore the relative similarity of TRF weights in response to the phonetic features: high, low, back and front. If these phonetic contrasts, related to articulatory tongue movements, are perceptually relevant and represented at the neural level, then the TRF weights in response to each of these phonetic categories should have consistently similar responses, while they would be consistently different across categories.

## Statistical analysis

All statistical analyses directly comparing the groups were performed using repeated measures three-way ANOVAs. One-sample *t*-tests were used for post-hoc tests. Correction for multiple comparisons was applied where necessary via the Holm correction. In that case, the adjusted *p*-value was reported. Descriptive statistics for the neurophysiology results are reported as a combination of mean and standard error (SE).

## Data Availability

The consent given by participants at the outset of this study did not explicitly detail sharing of the data in any format. This limitation is in keeping with EU General Data Protection Regulation and is imposed by the Research Ethics Committees of the Capital Region of Denmark. These ethical restrictions prevent Eriksholm Research Centre from fully anonymising and sharing the dataset. To inquire on the possibility to access the data, please contact Claus Nielsen, Eriksholm research operations manager, at.

## Results

Behavioural scores indicated that participants were able to successfully perform the task, displaying a 73% correct performance on 2-choice questions regarding the target speech for the NR off condition, and an 84% correct performance in the NR on condition, with a significant impact of NR scheme on behavioural accuracy (Alickovic et al., 2021).

### Robust phonetic-feature TRF for target but not masker speech in HI listeners

Forward TRF models were fit to characterise the mapping between phonetic features and low-frequency (1-8 Hz) EEG signals. Separate TRF models were fit for target and masker speech streams, first with a phonetic-feature model F and then with a control model, F_*sh*_, where the phonetic categories were shuffled, while preserving their timing. A repeated-measures three-way ANOVA was conducted to evaluate the impact of NR (OFF and ON), model (F and F_*sh*_) and attention (target and masker) on the EEG prediction correlations. This analysis indicated a main effect of attention (*F*(1,33) = 24.50, *p* = 2.14e-05, ηp^2^ = 0.43), with larger EEG prediction correlations for the target compared to the masker speech (r_target_ > r_masker_, post-hoc *t*-test: *t*(33) = 4.95, *p* = 2.14e-05, Cohen’s *d* = 0.68); a main effect of model (*F*(1,33) = 31.62; *p* = 2.93e-06, ηp^2^ = 0.49), with the F model showing greater EEG prediction correlations than its control model, F_*sh*_, derived by re-running the TRF computation after shuffling the corresponding phoneme categories (r_F_ > r_Fsh_; post-hoc *t*-test: *t*(33) = 5.62, *p* = 2.93e-06, Cohen’s *d* = 0.54); and a significant Attention*Model interaction (*F*(1,33) = 18.82, *p* = 1.28e-04, ηp^2^ = 0.36). The interaction effect revealed that an increase in EEG prediction correlation in the F model compared to its shuffled version F_*sh*_ (rF > rF_*sh*_) emerged for the target stimulus (post-hoc *t*-test: *t*(33) = 7.10, *p* = 6.98e-09, Cohen’s *d* = 0.87) whereas this effect did not emerge for the masker speech (post-hoc *t*-test: *t*(33) = 1.77, *p* = 0.16, Cohen’s *d* = 0.21). Finally, we did not find a main effect of NR (*F*(1,33) = 0.17; *p* = 0.68, ηp^2^ = 0.005).

For simplicity, we only report plots of the EEG prediction correlations (averaged across all EEG channels) and single-channel TRF weights (**Figure 2B**) for the NR OFF condition. Statistical post-hoc tests showed that the F model yields significantly higher prediction scores than its shuffled F_*sh*_ version for the target (post-hoc *t*-test: *t*(33) = 5.11, *p* = 2.96e-05, Cohen’s *d* = 0.85), but not for the masker speech (post-hoc *t*-test: *t*(33) = 0.65, *p* = 1, Cohen’s *d* = 0.15). Furthermore, an attentional effect emerged, since EEG prediction correlation values were significantly greater for the target compared to the masker, for both the F model (post-hoc *t*-test: *t*(33) = 4.67, *p* = 1.96e-04, Cohen’s *d* = 0.1) and the F_*sh*_ model (post-hoc *t*-test: *t*(33) = 1.79, *p* = 1, Cohen’s *d* = 0.38); (**Figure 2A)**. Similarly, for the NR ON condition, the F model’s EEG prediction correlations were significantly greater than those of the control F_*sh*_ model in the case of the target stream (post-hoc *t*-test: *t*(33) = 5.41, *p* = 9.14e-06, Cohen’s *d* = 0.9), while this result did not emerge for the masker (post-hoc *t*-test: *t*(33) = 1.23, *p* = 1, Cohen’s *d* = 0.2); (**Figure 2-2A** in **Extended Data)**. As for the NR OFF condition, also in the NR ON the accuracy of the EEG prediction was overall greater for the target compared to the masker speech, but for the F model only (post-hoc *t*-test: *t*(33) = 4.8, *p* = 1.31e-04, Cohen’s *d* = 1), while in this portion of the data there was no difference between target and masker in the F_*sh*_ model (post-hoc *t*-test: *t*(33) = 1.54, *p* = 1, Cohen’s *d* = 0.33). TRF weights for channel FCz in the NR ON condition are visualised in **Figure 2-2B** in **Extended Data**.

**Figure 2.**
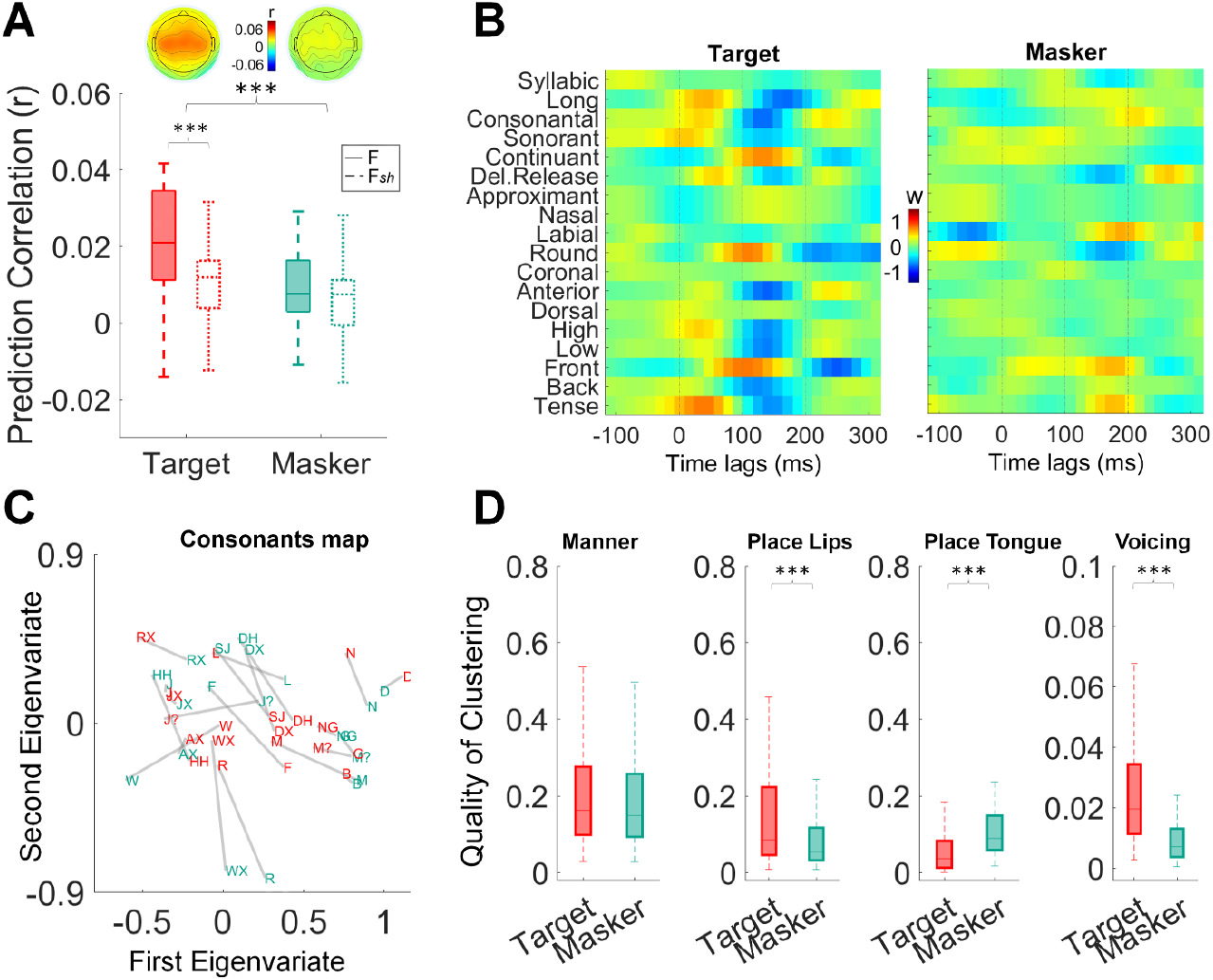
Stronger cortical encoding of target acoustic-phonetic information compared to masker information in listeners with hearing-impairment. **(A)** EEG prediction correlations (Pearson’s r) for phonetic features models F and shuffled control F_sh_ of the target and masker speech. Scalp topographies represent the distribution of prediction correlations across all channels (average across participants) for the F model. Error bars indicate the SEM across subjects. **(B)** TRF weights at channel FCz for the eighteen phonetic features for the F model. **(C)** Phoneme distance maps (PDMs) for target (red) and masker (green) speech. **(D)** EEG sensitivity to groups of phonetic features, i.e., quality of clustering of the EEG responses around relevant phonetic contrasts, for target and masker speech.

The results above indicate a robust relationship between EEG signals and acoustic-phonetic information of the target speech, while no such relationship is observed for the masker speech. Subsequent analyses were conducted to clarify whether attention had a more pronounced influence on the encoding of specific groups of phonetic features. To do so, PDMs reflecting the encoding of the target and masker speech were derived from the TRF weights of the target and masker F models separately (**Figure 2C**; see **Methods; Figure 2-2C** for NR ON in **Extended Data**). Calinski-Harabasz metrics were derived to evaluate the quality of clustering for both the target and masker speech. These metrics measure the degree of separation between the relevant phonetic feature clusters within the EEG-based PDMs. Larger values indicate that the organisation of PDMs aligns well with a specific group of features.

**Figure 2D** summarises the results of the PDM analysis indicating that, overall, the neural encoding of different phonetic feature sets is more consistently clustered in the target compared to the masker speech, while the impact of NR emerged as significant for some phonetic features sets, but not for others. These results emerged from a two-way repeated measures ANOVA – NR (ON and OFF) and attention (target and masker) - which was conducted four times, one for each phonetic feature set.

For *manner of articulation*, the analysis revealed a main effect of attention (*F*(1,99) = 15.90; *p* = 1.29e-04, ηp^2^ = 0.14), with a general effect of target > masker, (post-hoc *t*-test: *t*(33) = 3.99, *p* = 1.29e-04, Cohen’s *d* = 0.36), while there was no effect of NR (*F*(1,99) = 2.79; *p* =0.09, ηp^2^ = 0.03). In the specific case of NR OFF plotted here, there was no statistical difference between target and masker (post-hoc *t*-test: *t*(33) = 1.74, *p* = 0.25, Cohen’s *d* = 0.23). For the NR ON condition (**Figure 2-2D** in **Extended Data**), the clustering of *manner*-related phonetic features was more consistent for the target speech compared to the masker (post-hoc *t*-test: *t*(33) = 3.8, *p* = 9.61e-04, Cohen’s *d* = 0.5).

For *place of articulation (lips)*, a significant effect of attention emerged (*F*(1,99) = 59.55; *p* = 9.65e-12, ηp^2^ = 0.38), with target > masker, (post-hoc *t*-test: *t*(33) = 7.72, *p* = 9.65e-12, Cohen’s *d* = 0.78). A main effect of NR was also found (*F*(1,99) = 12.47; *p* = 6.31e-4, ηp^2^ = 0.11), with NR ON > OFF (post-hoc *t*-test: *t*(33) = 3.53, *p* = 6.31e-4, Cohen’s *d* = 0.37), together with a statistically significant interaction of attention and NR (*F*(1,99) = 5.62; *p* = 0.02, ηp^2^ = 0.05). For the NR OFF condition, plotted in **Figure 2D**, the clustering for the target speech was greater than that for the masker (post-hoc *t*-test: *t*(33) = 4.10, *p* = 1.82e-4, Cohen’s *d* = 0.56). The same pattern of results emerged in the NR ON condition, plotted in **Figure 2-2D** in **Extended Data** (post-hoc *t*-test: *t*(33) = 7.3, *p* = 3.57e-11, Cohen’s *d* = 1).

Regarding the *place of articulation (tongue)* features, the ANOVA revealed once again a significant main effect of attention (*F*(1,99) = 45.86; *p* = 9.15e-10, ηp^2^ = 0.32), this time with a more effective clustering for the masker than for the target, (post-hoc *t*-test: *t*(33) = 6.77, *p* = 9.15e-10, Cohen’s *d* = 0.64). For NR OFF specifically, it can be observed that masker > target (post-hoc *t*-test: *t*(33) = 3.50, *p* = 0.002, Cohen’s *d* = 0.50). Similarly, in the case of the NR ON condition (**Figure 2-2D** in **Extended Data**), the masker speech showed a more consistent clustering than the target (post-hoc *t*-test: *t*(33) = 5.4, *p* = 1.19e-06, Cohen’s *d* = 0.79).

In the case of voicing, the 2-way ANOVA revealed a significant effect of attention (*F*(1,99) = 221.96; *p* = 4.96e-27, ηp^2^ = 0.70), with target > masker (post-hoc *t*-test: *t*(33) = 14.90, *p* = 4.96e-27, Cohen’s *d* = 1.46), a significant effect of NR (*F*(1,99) = 14.35; *p* = 2.61e-4, ηp^2^ = 0.13), with NR ON yielding a better clustering than NR OFF (post-hoc *t*-test: *t*(33) = 3.80, *p* = 2.61e-4, Cohen’s *d* = 0.39), and a significant attention*NR interaction (*F*(1,99) = 48.23; *p* = 4.03e-10, ηp^2^ = 0.33). For the NR OFF condition reported here, phonetic features related to voicing were more consistently clustered for the target compared to the masker speech (post-hoc *t*-test: *t*(33) = 6.54, *p* = 1.03e-9, Cohen’s *d* = 0.86). Similarly, the NR ON condition (**Figure 2-2D** in **Extended Data**) showed a significantly greater clustering for the target compared to the masker stream (post-hoc *t*-test: *t*(33) = 15.76, *p* = 7.54e-36, Cohen’s *d* = 2.07).

### Cortical encoding of both target and masker phoneme onsets in HI listeners

The TRF F model potentially captured both an acoustically invariant cortical encoding of phonetic features as well as responses to other correlated information, such as the acoustic spectrogram (Teoh et al., 2022). To isolate EEG signals that can be explained by phonetic feature categories but not by speech acoustics, TRF models were fit based on an FS set of features comprising the S model (spectrogram and the half-wave rectified spectrogram derivative) and the F model (phonetic-feature categories). The gain in EEG prediction correlations due solely to phonetic features was then assessed by subtracting the FS and the S models, yielding the PhCat metric.

The PhCat metric (FS-S gain) may reflect the EEG encoding of the timing of phonetic feature categories, as well as their identity (Di Liberto et al., 2015; Di Liberto et al., 2021; Liberto et al., 2022). To assess if either or both these properties were encoded in the EEG signals, we also compared the FS-S predictive performance with the results of the same analysis after shuffling the phoneme identities in the transcript, which resulted in the PhOnset metric (F_*sh*_S_-_S gain). This PhOnset metric allowed us to isolate the contribution of phonetic feature onsets, while the PhCat metric represented the contribution of phonetic feature identity.

We compared the EEG prediction correlation gain obtained from the PhCat metric (FS-S) and the one obtained from the PhOnset metric (F_*sh*_*S*_*-*_*S*) using a repeated-measures three-way ANOVA with factors: NR (OFF and ON), model gain (FS-S and F_*sh*_S-S) and attention (target and masker). This analysis was done to dissociate the potential contribution of phonetic-feature onsets from that of phonetic feature identity. Our analysis revealed no significant difference between the EEG prediction increase of the two model gains (PhCat and PhOnset metrics), suggesting that the contribution of phonetic feature onsets and phonetic feature identity is overall comparable (no main effect of model gains: *F(*1,33) = 1.98, *p* = 0.17, ηp^2^ = 0.06). Furthermore, we observed a significantly greater EEG prediction increase for the masker speech, compared to the target, across the two model gains (significant main effect of attention: *F(*1,33) = 11.26, *p* = 0.002, ηp^2^ = 0.25. r_masker_ > r_target_, post-hoc *t*-test: *t*(33) = 3.36, *p* = 0.002, Cohen’s *d* = 0.42). The ANOVA results did not indicate a significant main effect of NR on the EEG prediction correlations (*F(*1,33) = 0.09, *p* = 0.7, ηp^2^ = 0.003). An interaction between attention and model gain did reach statistical significance (*F(*1,33) = 4.26, *p* = 0.047, ηp^2^ = 0.11), showing that the target speech PhOnset metric yielded significantly lower EEG prediction gains compared to both the masker PhCat metric (post-hoc *t*-test: *t*(33) = -3.63, *p* = 0.003, Cohen’s *d* = -0.5) and the masker PhOnset metric (post-hoc *t*-test: *t*(33) = -3.94, *p* = 0.001, Cohen’s *d* = -0.56).

As in the previous paragraph, we only report plots for the NR OFF condition, where no difference was found between the PhCat and PhOnset metrics, neither for the target speech (post-hoc *t*-test: *t*(33) = 1.10, *p* = 1, Cohen’s *d* = 0.15) nor for the masker speech (post-hoc *t*-test: *t*(33) = 1.05, *p* = 1, Cohen’s *d* = 0.14). The statistical analysis showed a greater predictive gain of phoneme onsets for the masker speech, compared to the target (post-hoc *t*-test: *t*(33) = 3.24, *p* = 0.046, Cohen’s *d* = 0.72) (**Figure 3A and 3B)**. For the NR ON condition, we found the same pattern of results, with no significant difference between the PhCat and PhOnset metrics, neither for the target (post-hoc *t*-test: *t*(33) = 2.38, *p* = 0.43, Cohen’s *d* = 0.32) nor for the masker stream (post-hoc *t*-test: *t*(33) = 1.86, *p* = 1, Cohen’s *d* = 0.41) (**Figure 3-2A and 3-2B** in **Extended Data)**.

**Figure 3.**
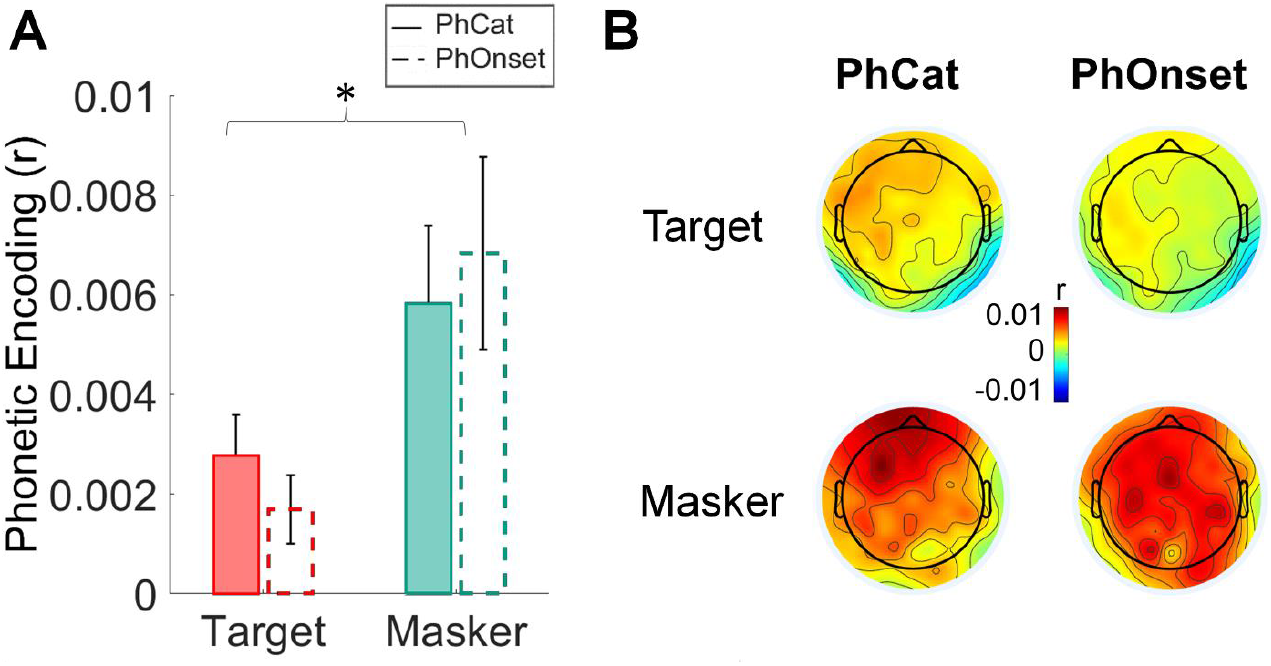
Over-representation of phonemic onsets of the ignored speech sounds in the low-frequency EEG of HI participants. **(A)** EEG prediction gains obtained from the PhCat (FS-S) and PhOnset (F_sh_S-S) metric, for the target and masker speech. Bars represent the increase in prediction correlations (r) averaged across all subjects and electrodes. Error bars represent the SEM across subjects. **(B)** Topographical distribution of the average EEG prediction correlation increases from the baseline model S, across all electrode locations.

## Discussion

Individuals with hearing impairment have difficulties in sustaining attention to the speaker of interest in multi-talker scenarios (Lunner et al., 2020). Here, we measured how attentional selection modulates the cortical encoding of phonetic features in HI individuals, testing whether hearing loss leads to an over-representation of the masker speech. Our results show two main patterns: First, we found that cortical responses in HI participants relate to phonetic features more strongly for the target than the masker streams, which is similar to the pattern previously measured in NH individuals (Teoh et al., 2022). Second, when isolating the different contributors to phonetic feature encoding, we found a significant cortical encoding of phoneme onsets of both target and masker speech, with stronger a representation of the masker.

Our findings impact the current understanding of how selective attention strategies unfold in the brain of HI listeners. The finding of a stronger relationship between cortical responses and phonetic features suggests that HI individuals perform attentional filtering at the level of the phonetic-feature encoding, segregating target and masker stream and processing phonetic information of target and masker in a different manner. However, we could not isolate the acoustically invariant component of the phonetic responses here, meaning that the effect of selective attention on the phonetic feature EEG metric likely reflects a combination of acoustically variant and invariant responses to phonetic features. While possible explanations could relate to sample size and data quality, it should be noted that this is the first study measuring phonetic feature TRFs with the Danish language, while previous studies isolating acoustically invariant phonetic-feature TRFs used the English language (and Flemish), (Gillis et al., 2023). Future work could further investigate this question by considering NH and HI participants across different languages.

Our finding that cortex of HI individuals encodes the phoneme onsets of both target and masker speech confirms the initial hypothesis of an over-representation of phoneme-level information of the masker, which could explain the high distractibility reported by the participants. Interestingly, the cortical encoding of phoneme onsets was stronger for the masker than the target. This result suggests that the onset encoding itself changes with the specific phoneme category for the target speech, reducing the consistency across different occurrences when those categories are shuffled in F_*sh*_. Conversely, the stronger phoneme onset encoding for the masker suggests higher consistency across phoneme categories, as it emerged from the analysis in **Fig. 2A**. Overall, these results indicate a significant encoding of phonetic distinctions for the target but not the masker, as well as an over-representation of the masker phoneme onsets. In the context of selective attention, the cortical tracking of the masker phoneme onsets could serve two important functions: keeping a fundamental structural representation of the to-be-ignored stimulus for an effective re-orienting of attention on the masker stimulus at later stages and building a structural representation of the masker speech, which is useful for suppressing it.

It is interesting to note that the EEG-measured phoneme-level encoding was not influenced by the use of a hearing-aid noise-reduction scheme, despite increased comprehension. The noise-reduction scheme reduced the sound level of the 16-talker babble noise in the background, without affecting the relative sound level between the target and masker speech streams originating in front of the speaker. As such, while NR ON made the task easier increasing the overall comprehension (Alickovic et al., 2020; Alickovic et al., 2021), the listener’s brain still had to segregate target and masker speech streams. Based on this result, we can speculate that the impact of attentional modulation measured at the phonetic level does not reflect overall speech comprehension, but a more general cortical strategy for selective attention in HI participants. Instead, measuring overall speech comprehension might be effective by isolating higher order processes, such as lexical predictions (Gillis et al., 2021).

Previous work highlighted the key role of temporal information and phoneme onsets in the segregation of auditory streams in multi-talker scenarios. Temporal coherence was proposed as a key criterion for grouping acoustic information into separate streams (Elhilali et al., 2009; O’Sullivan et al., 2015a). Previous work also found acoustic onsets of naturalistic speech to be faithfully represented in the auditory cortex (Hamilton et al., 2018; Daube et al., 2019), guiding the segregation of target and masker streams (Brodbeck et al., 2020). Acoustic-onset enhancement of the target speech signal has proven to benefit sentence recognition in cochlear implant users, even in the presence of competing speech (Stilp and Kluender, 2010; Koning and Wouters, 2012). Furthermore, the neural encoding of acoustic onsets of the masker speech streams was suggested to serve as a template structure of the masker stream, contributing to its suppression (Brodbeck et al., 2020). Intriguingly, the late encoding of acoustic onsets of the masker in fronto-parietal regions was also shown to be important for attentional selection, with a stronger encoding emerging when the masker stream was presented at a higher sound level than the target stream (Fiedler et al., 2019). This effect could not be studied with our experimental paradigm, which only modulated the SNR by reducing the sound level of the speech babble in the background (NR OFF vs. NR ON), not the relative sound-level of target and masker streams. Further work exploring different relative sound-levels could inform us on whether the late encoding of acoustic onsets measured in fronto-parietal areas is purely acoustic or related to phonological processing. Future investigations could also investigate the relationship between age and the degree of hearing loss with the effects measured in the present study, further explaining what the over-representation of phonetic onsets reflects exactly.

In conclusion, this study presented novel insight into the impact of selective attention on the neural encoding of speech in listeners with hearing loss. Our results complement previous work on attentional selection by measuring phonological neural encoding in a naturalistic multi-talker scenario. The finding that target and masker streams are segregated in the human cortex is in line with models of object-based auditory attention (Shinn-Cunningham, 2008; Shinn-Cunningham and Best, 2008) and complements previous evidence on the encoding of masker speech streams (Brodbeck et al., 2020), informing us for the first time on the neural underpinnings of this process at the phoneme level in HI participants.

## Author Contributions

Neural data were collected by EA and research clinicians as part of a previous study. The study was conceived by EA, GDL, ALV, and JZ.

Data were re-analysed by SC under the supervision of GDL, ALV, EA, and JZ. SC and GDL wrote the first draft of the manuscript.

JZ, EA, ALV edited the manuscript.

## Acknowledgements

This work was conducted with the financial support of the William Demant Foundation, grant 21-0628 and grant 22-0552 and of the Science Foundation Ireland Centre for Research Training in Artificial Intelligence, under Grant No. 18/CRT/6223. This research was supported by the Science Foundation Ireland under Grant Agreement No. <u>13/RC/2106_P2</u> at the ADAPT SFI Research Centre at Trinity College Dublin. ADAPT, the SFI Research Centre for AI-Driven Digital Content Technology, is funded by Science Foundation Ireland through the SFI Research Centres Programme.

