## Extended Data for "Cortical over-representation of phonetic onsets of ignored speech in hearing impaired individuals"

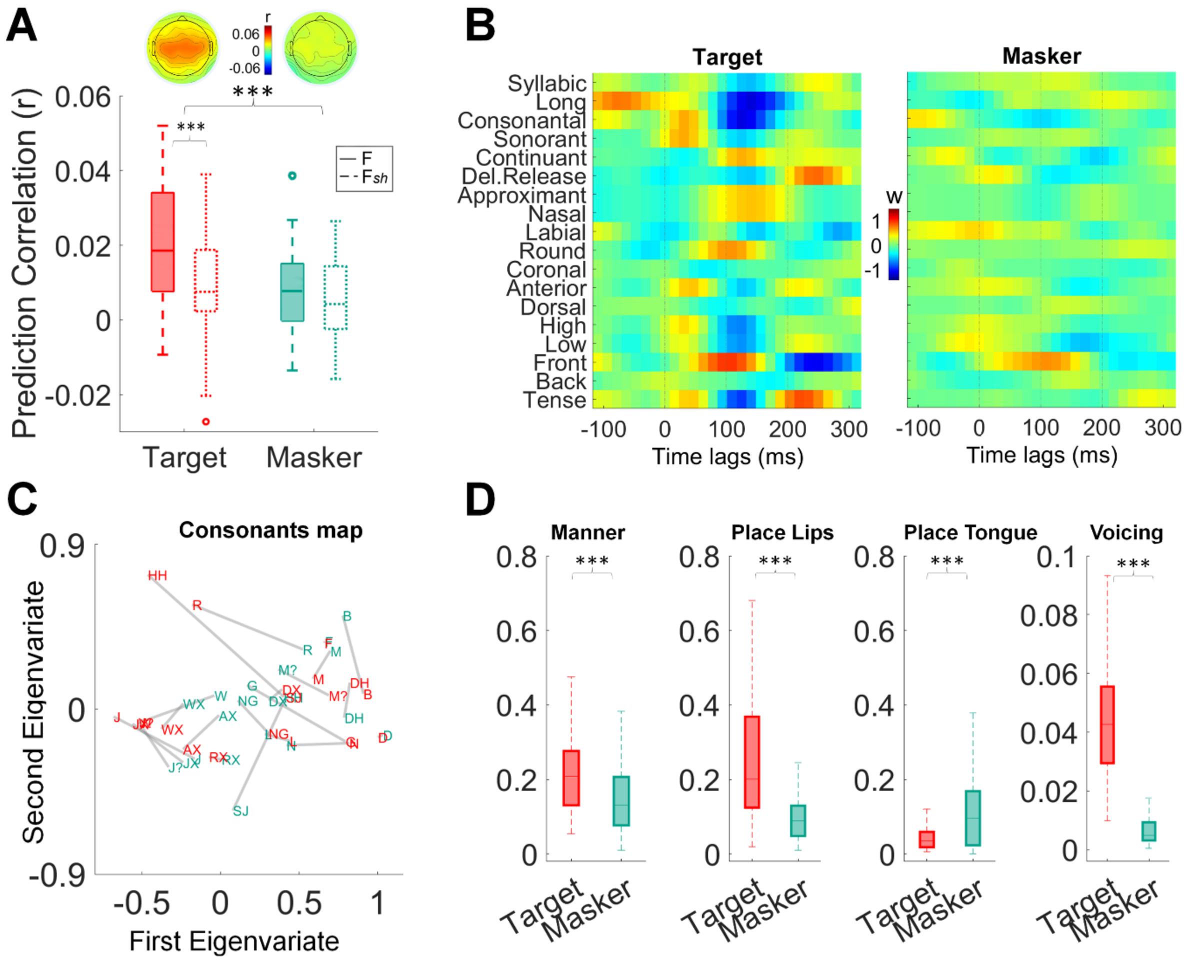


***Figure 1-2. Target acoustic-phonetic information is more strongly encoded than the masker's information in listeners with hearing impairment when background noise is removed in the NR ON condition. (A****) EEG prediction correlations (Pearson's r) for phonetic features models F and shuffled control $F_{sh}$ of the target and masker speech. Scalp topographies represent the distribution of prediction correlations across all channels (average across participants) for the F model. Error bars indicate the SEM across participants.* ***(B)*** *TRF weights at channel FCz for the eighteen phonetic features for the F model.* ***(C)*** *Phoneme distance maps (PDMs) for target (red) and masker (green) speech.* ***(D)*** *EEG sensitivity to groups of phonetic features, i.e., quality of clustering of the EEG responses around relevant phonetic contrasts, for target and masker speech.*

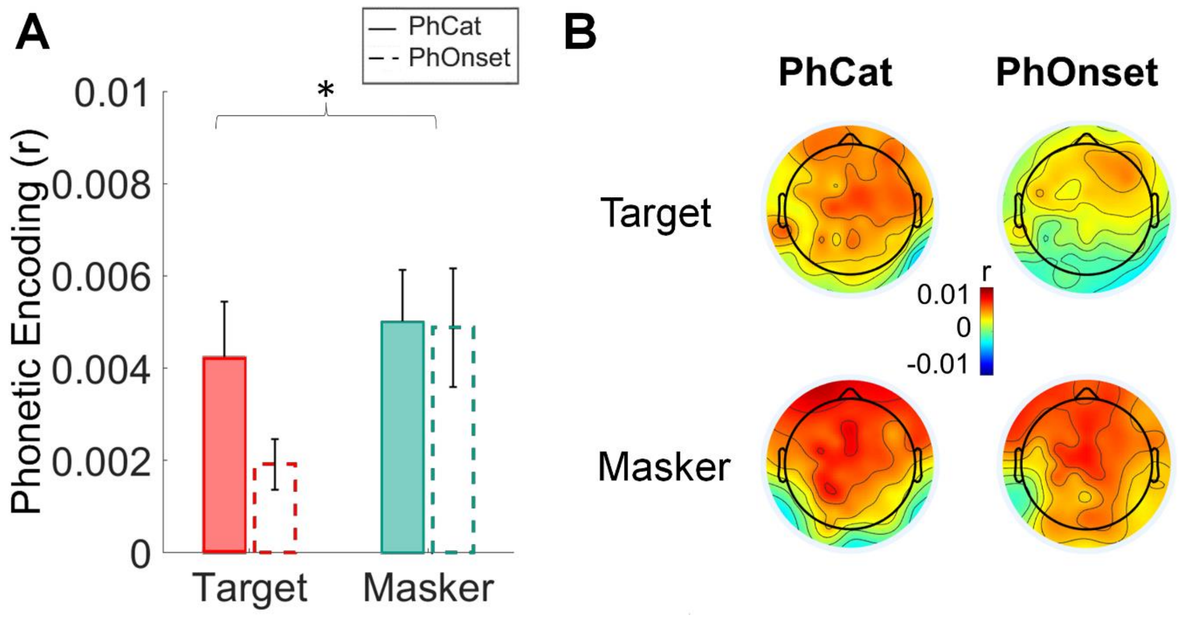


***Figure 3-2. Over-representation of phonemic onsets of the ignored speech sounds in the low-frequency EEG of hearing-impaired participants when background babble noise is suppressed in the NR ON condition. (A)*** *EEG prediction gains obtained from the PhCat (FS-S) and PhOnset ($F_{sh}$S-S) metric, for the target and masker speech. Bars represent the increase in prediction correlations (r) averaged across all subjects and electrodes. Error bars represent the SEM across subjects.* ***(B)*** *Topographical distribution of the average EEG prediction correlation increases from the baseline model S, across all electrode locations.*
